## Supplementary Figures for "Cerebrospinal fluid metabolomics identifies 19 brain-related phenotype associations"

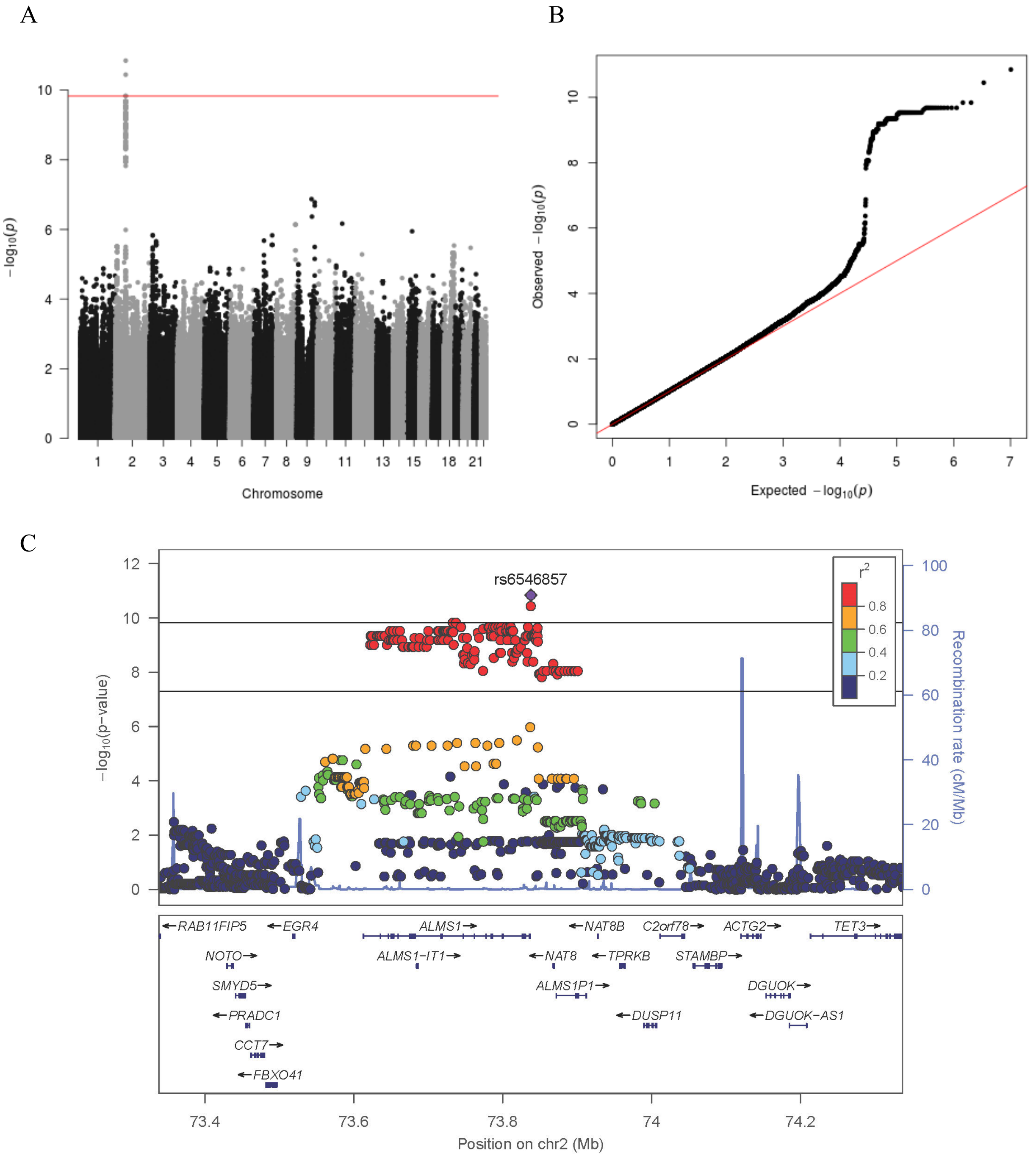

**Supplementary Figure 1. GWAS meta-analysis results for methionine sulfone (X44748)**

Panel A shows the Manhattan plot, with a red line indicating the Bonferroni-corrected significance threshold (P = $1.48 \times 10^{-10}$). Panel B shows the Q-Q plot of the estimated effect sizes. Panel C shows the regional association plot for the associated locus. The top horizontal line represents the Bonferroni-corrected p-value threshold while the bottom line represents the standard genome-wide significance threshold.

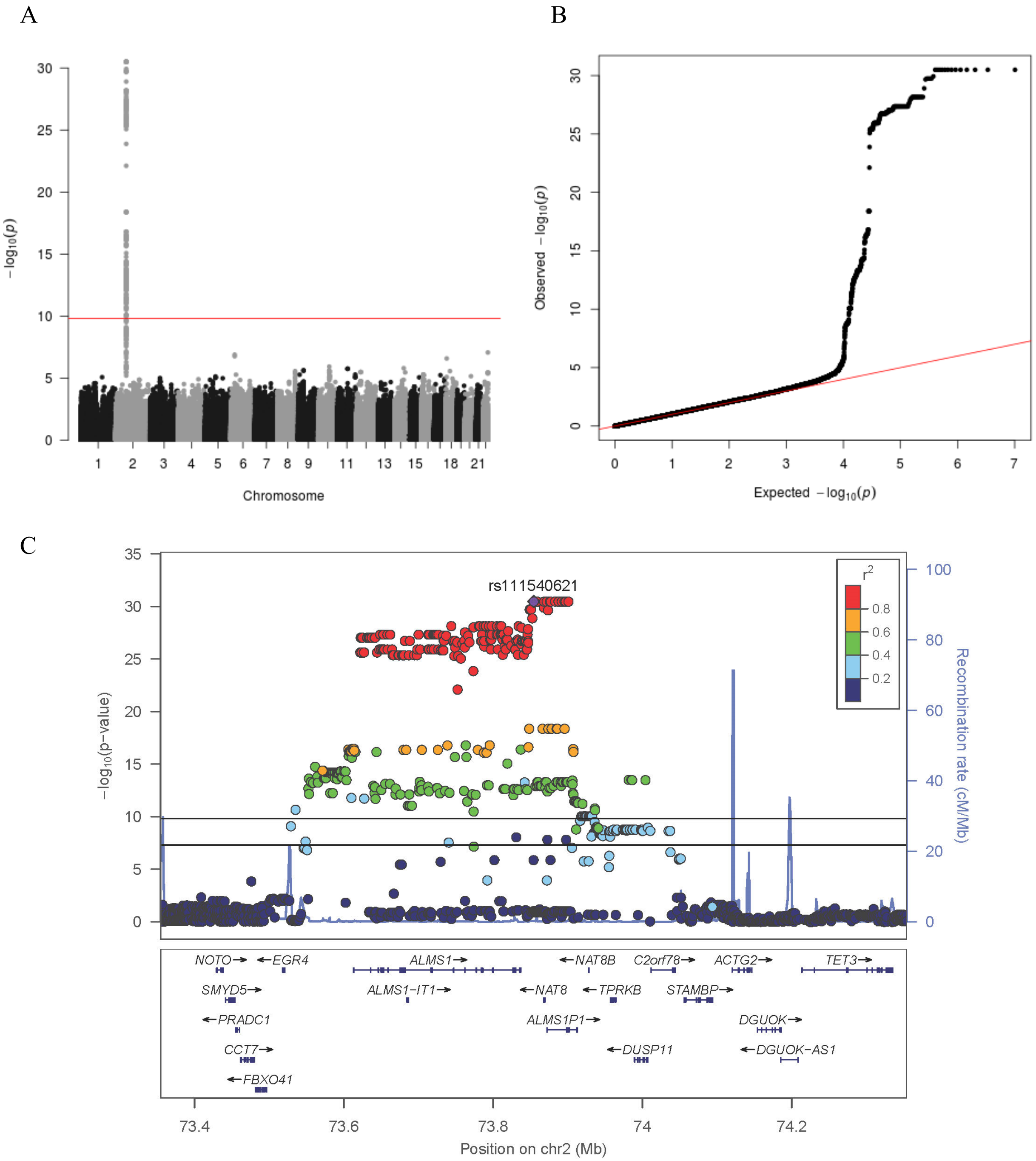

**Supplementary Figure 2. GWAS meta-analysis results for N-delta-acetylornithine (X43249)**

Panel A shows the Manhattan plot, with a red line indicating the Bonferroni-corrected significance threshold ($P = 1.48x10^{-10}$). Panel B shows the Q-Q plot of the estimated effect sizes. Panel C shows the regional association plot for the associated locus. The top horizontal line represents the Bonferroni-corrected p-value threshold while the bottom line represents the standard genome-wide significance threshold.

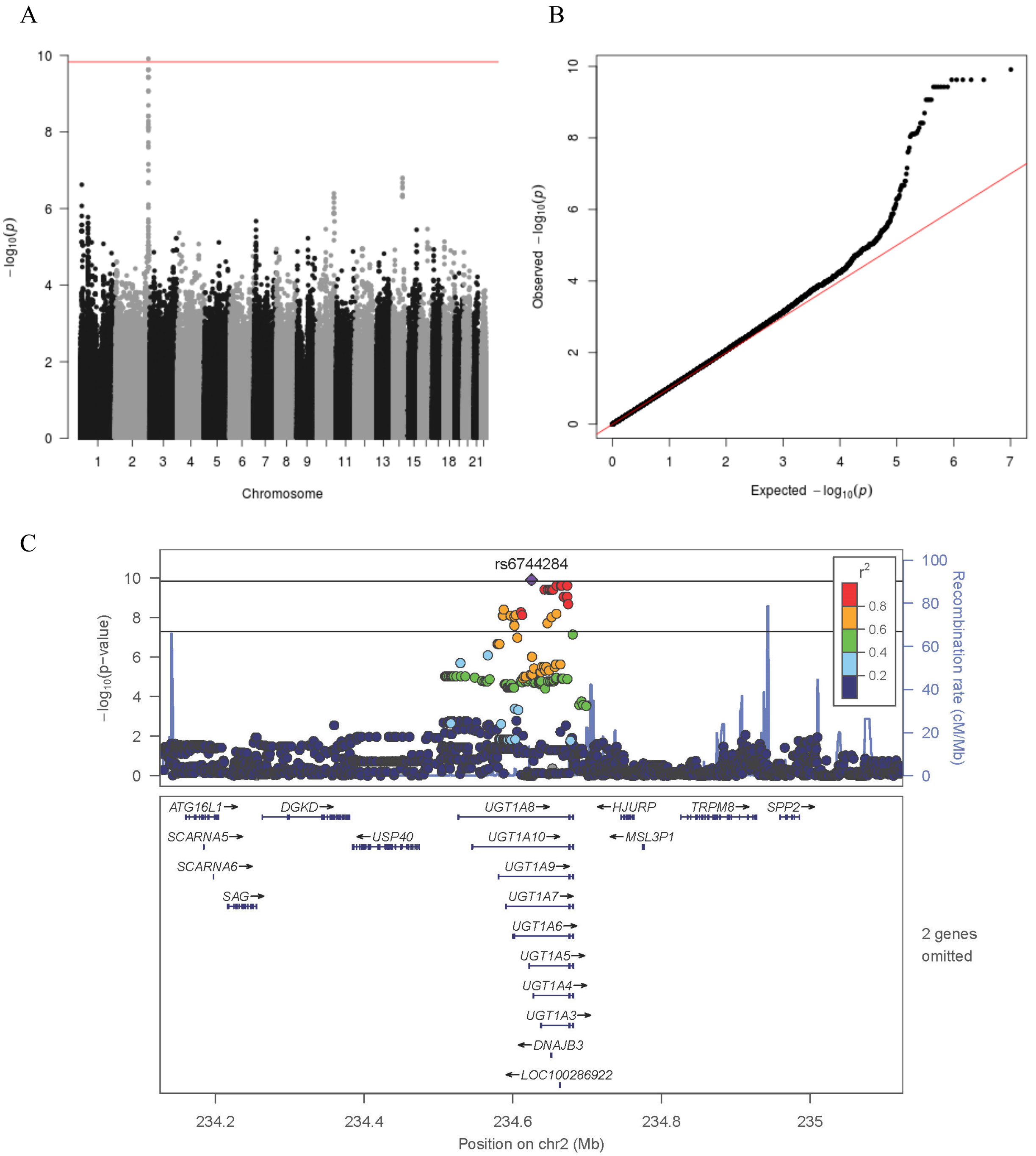

**Supplementary Figure 3. GWAS meta-analysis results for bilirubin (X43807)**

Panel A shows the Manhattan plot, with a red line indicating the Bonferroni-corrected significance threshold ($P = 1.48x10^{-10}$). Panel B shows the Q-Q plot of the estimated effect sizes. Panel C shows the regional association plot for the associated locus. The top horizontal line represents the Bonferroni-corrected p-value threshold while the bottom line represents the standard genome-wide significance threshold.

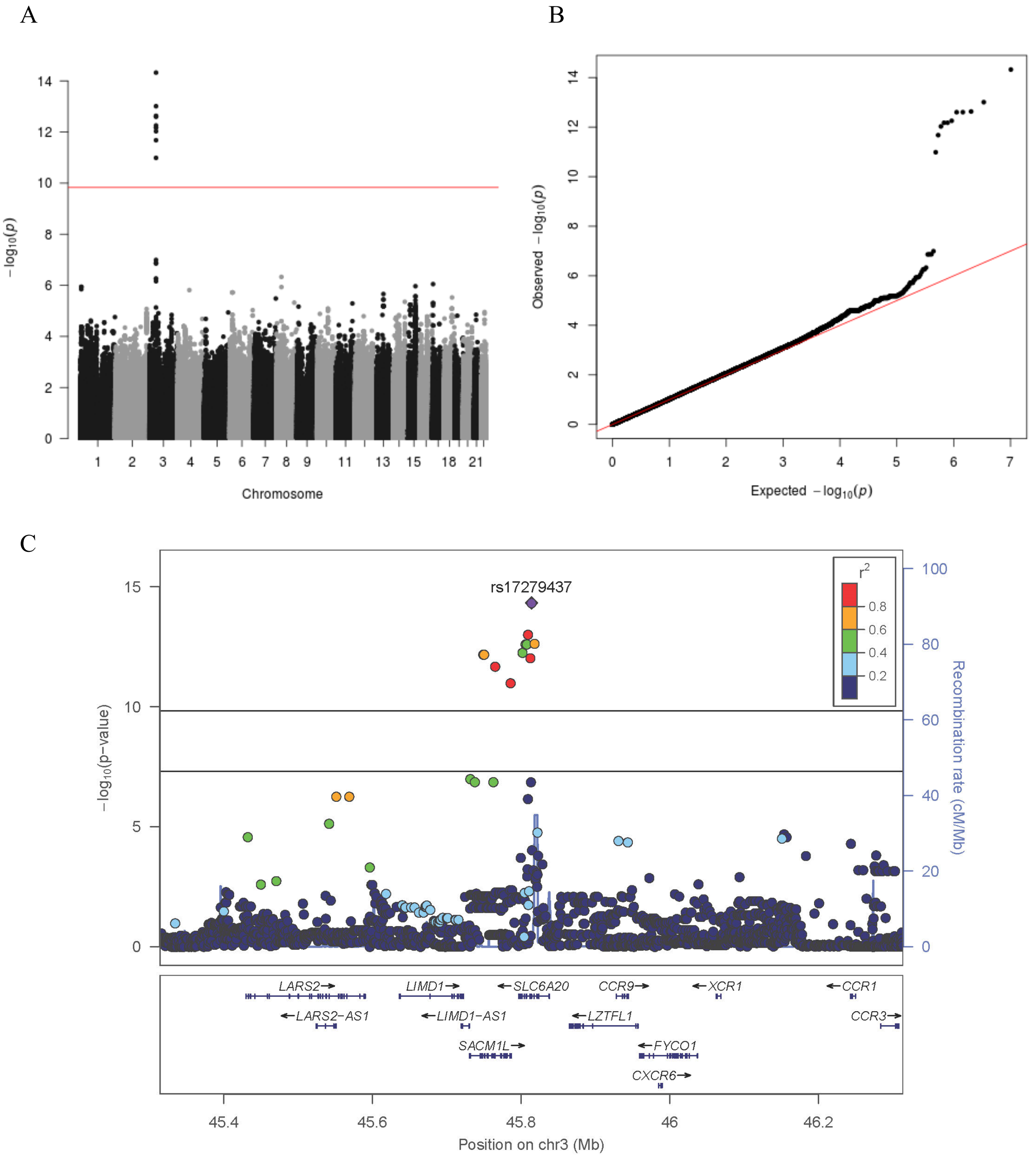

**Supplementary Figure 4. GWAS meta-analysis results for betaine (X3141)**

Panel A shows the Manhattan plot, with a red line indicating the Bonferroni-corrected significance threshold (P = $1.48 \times 10^{-10}$). Panel B shows the Q-Q plot of the estimated effect sizes. Panel C shows the regional association plot for the associated locus. The top horizontal line represents the Bonferroni-corrected p-value threshold while the bottom line represents the standard genome-wide significance threshold.

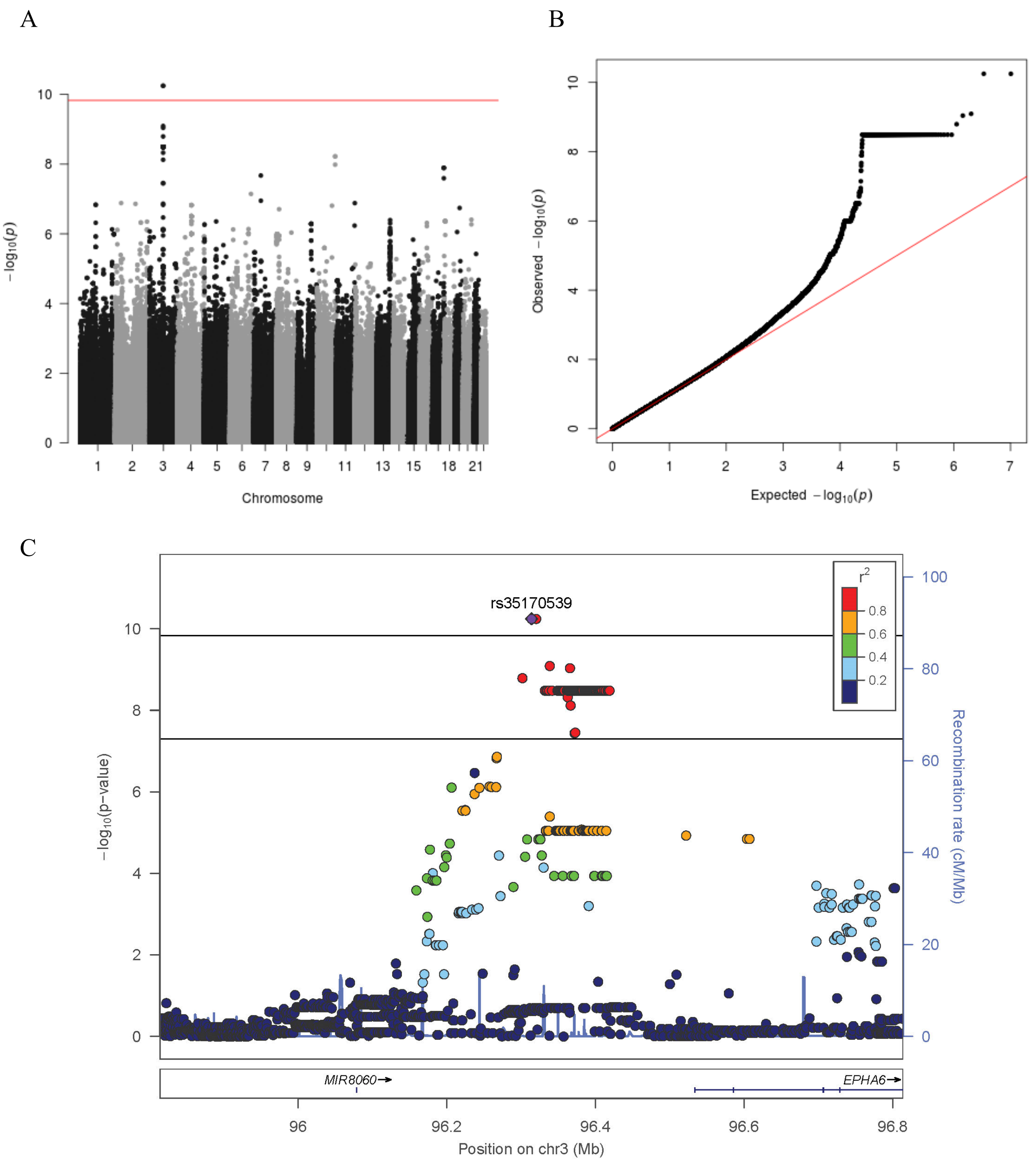

**Supplementary Figure 5. GWAS meta-analysis results for oxalate (ethanedioate) (X20694)**

Panel A shows the Manhattan plot, with a red line indicating the Bonferroni-corrected significance threshold ($P = 1.48 \times 10^{-10}$). Panel B shows the Q-Q plot of the estimated effect sizes. Panel C shows the regional association plot for the associated locus. The top horizontal line represents the Bonferroni-corrected p-value threshold while the bottom line represents the standard genome-wide significance threshold.

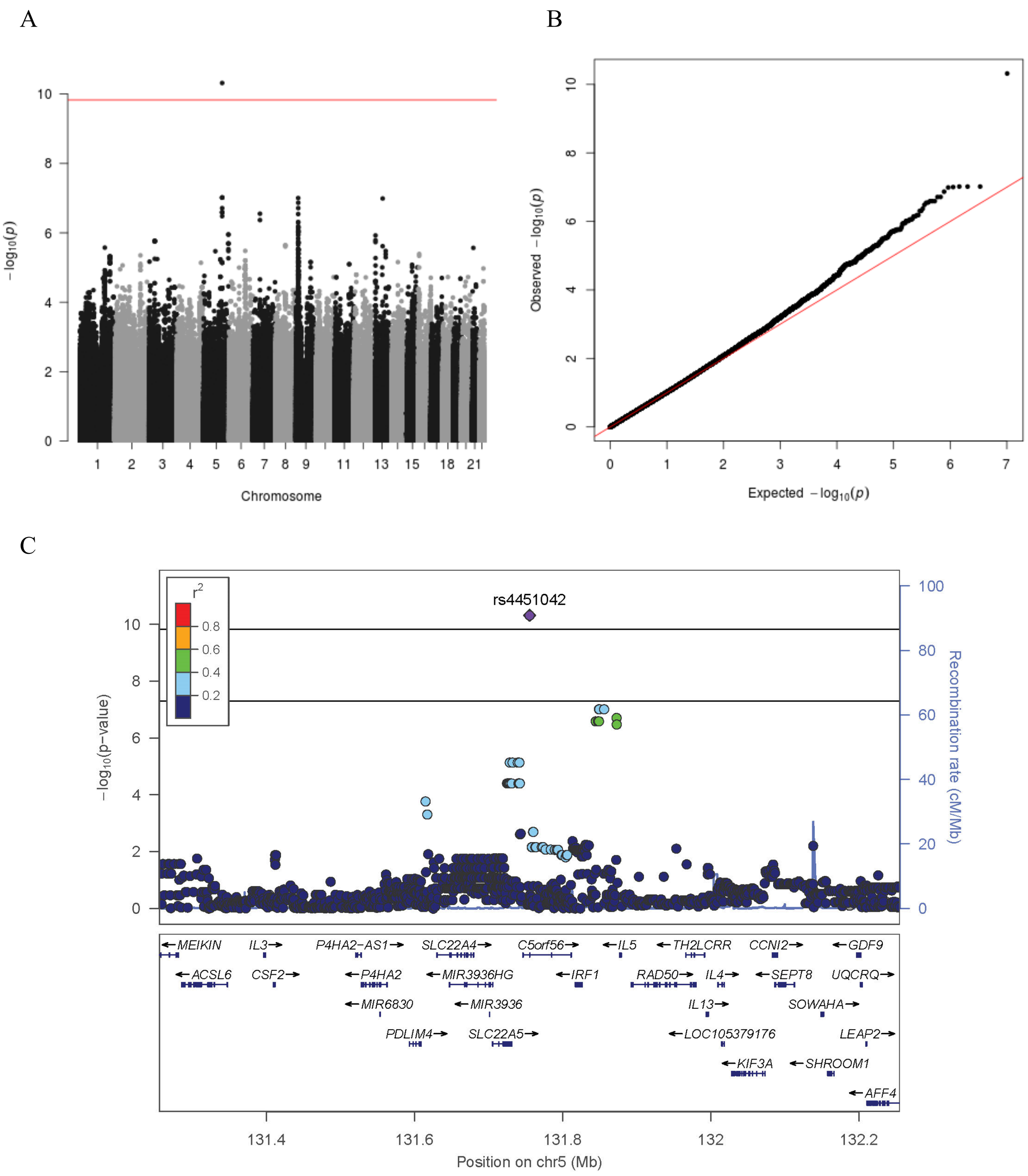

**Supplementary Figure 6. GWAS meta-analysis results for tryptophan betaine (X37097)**

Panel A shows the Manhattan plot, with a red line indicating the Bonferroni-corrected significance threshold (P = $1.48 \times 10^{-10}$). Panel B shows the Q-Q plot of the estimated effect sizes. Panel C shows the regional association plot for the associated locus. The top horizontal line represents the Bonferroni-corrected p-value threshold while the bottom line represents the standard genome-wide significance threshold.

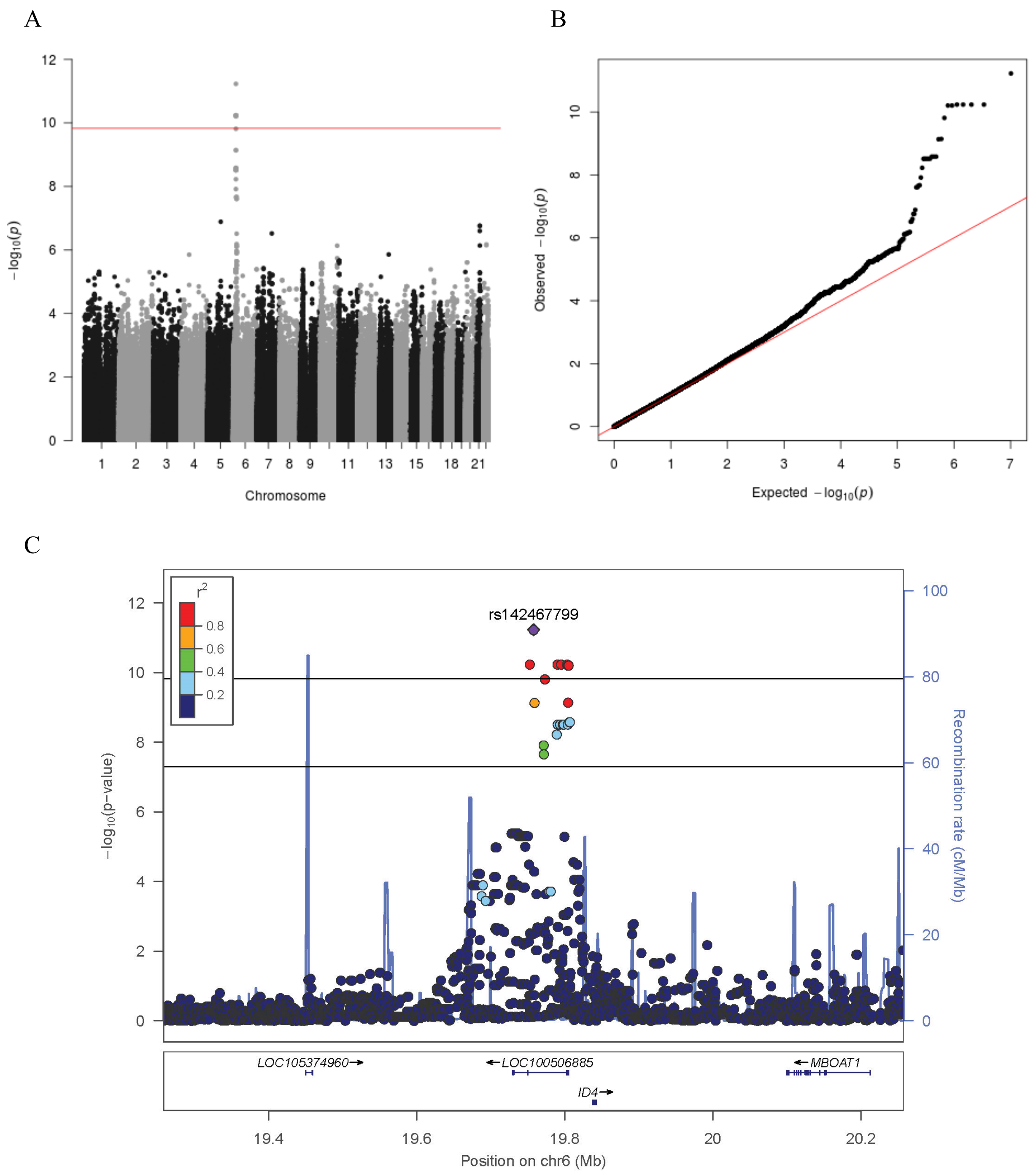

**Supplementary Figure 7. GWAS meta-analysis results for 2-hydroxyadipate (X31934)**

Panel A shows the Manhattan plot, with a red line indicating the Bonferroni-corrected significance threshold ($P = 1.48 \times 10^{-10}$). Panel B shows the Q-Q plot of the estimated effect sizes. Panel C shows the regional association plot for the associated locus. The top horizontal line represents the Bonferroni-corrected p-value threshold while the bottom line represents the standard genome-wide significance threshold.

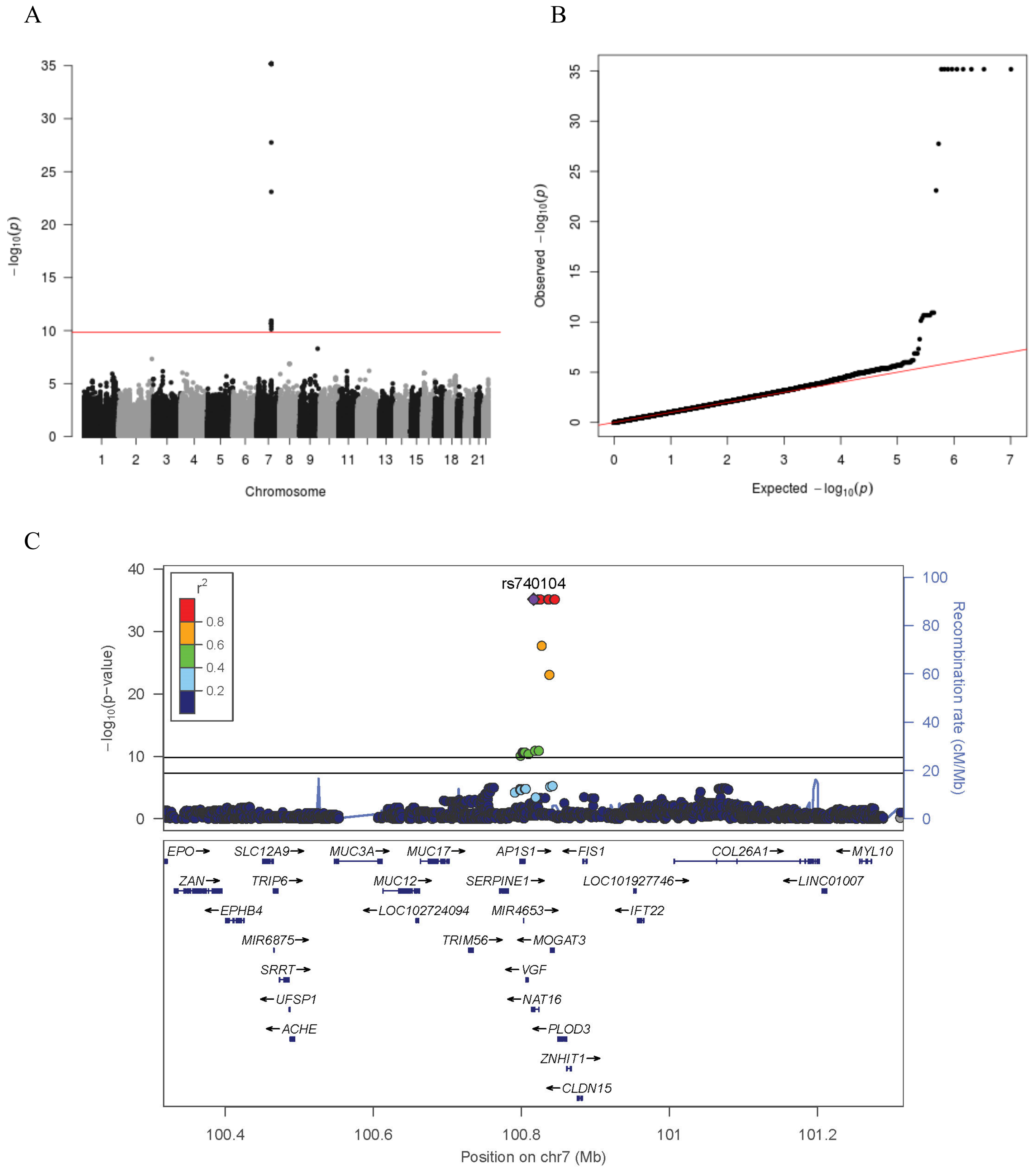

**Supplementary Figure 8. GWAS meta-analysis results for N-acetylhistidine (X33946)**

Panel A shows the Manhattan plot, with a red line indicating the Bonferroni-corrected significance threshold ($P = 1.48x10^{-10}$). Panel B shows the Q-Q plot of the estimated effect sizes. Panel C shows the regional association plot for the associated locus. The top horizontal line represents the Bonferroni-corrected p-value threshold while the bottom line represents the standard genome-wide significance threshold.

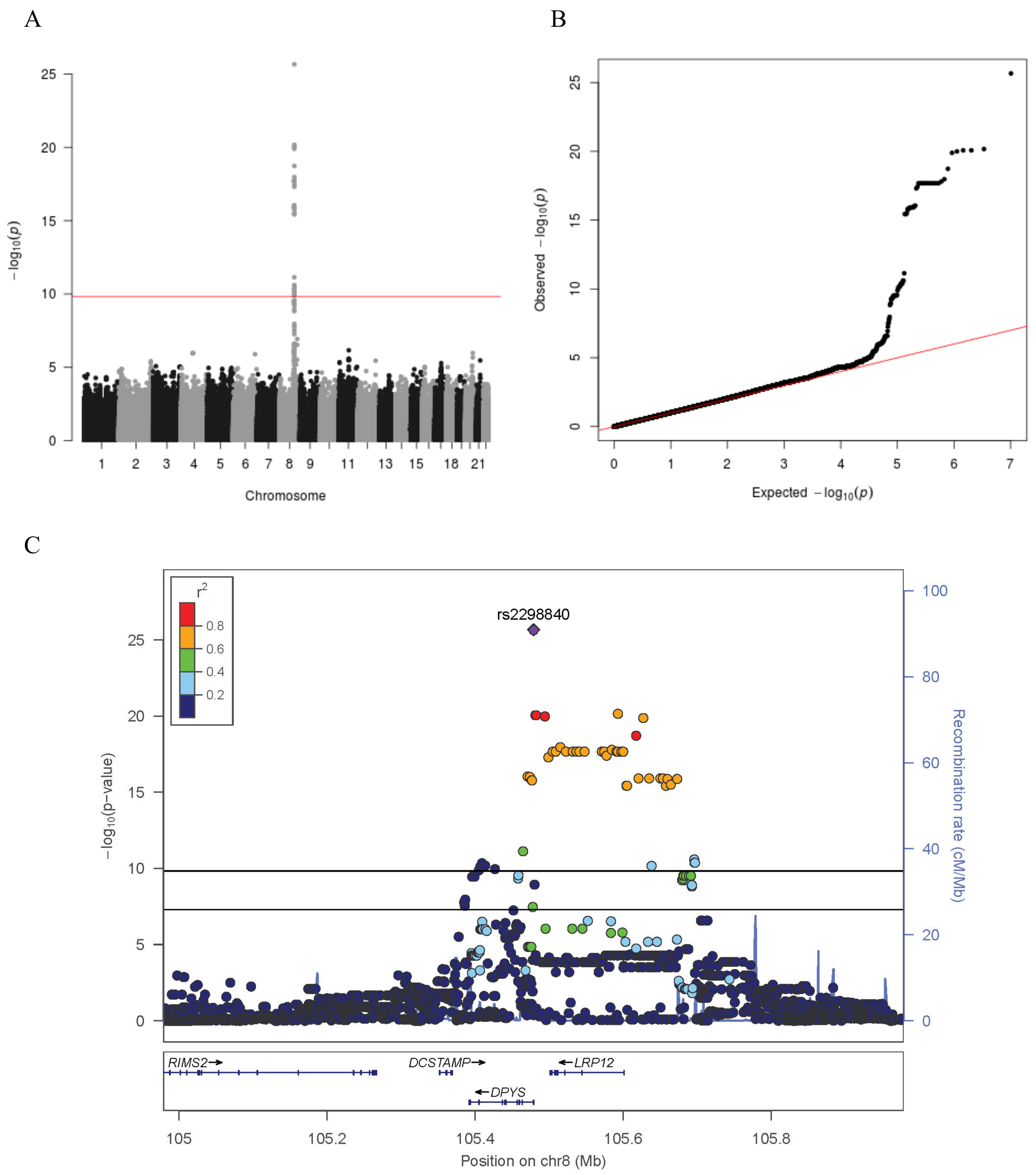

**Supplementary Figure 9. GWAS meta-analysis results for 3-ureidopropionate (X3155)**

Panel A shows the Manhattan plot, with a red line indicating the Bonferroni-corrected significance threshold (P = $1.48 \times 10^{-10}$). Panel B shows the Q-Q plot of the estimated effect sizes. Panel C shows the regional association plot for the associated locus. The top horizontal line represents the Bonferroni-corrected p-value threshold while the bottom line represents the standard genome-wide significance threshold.

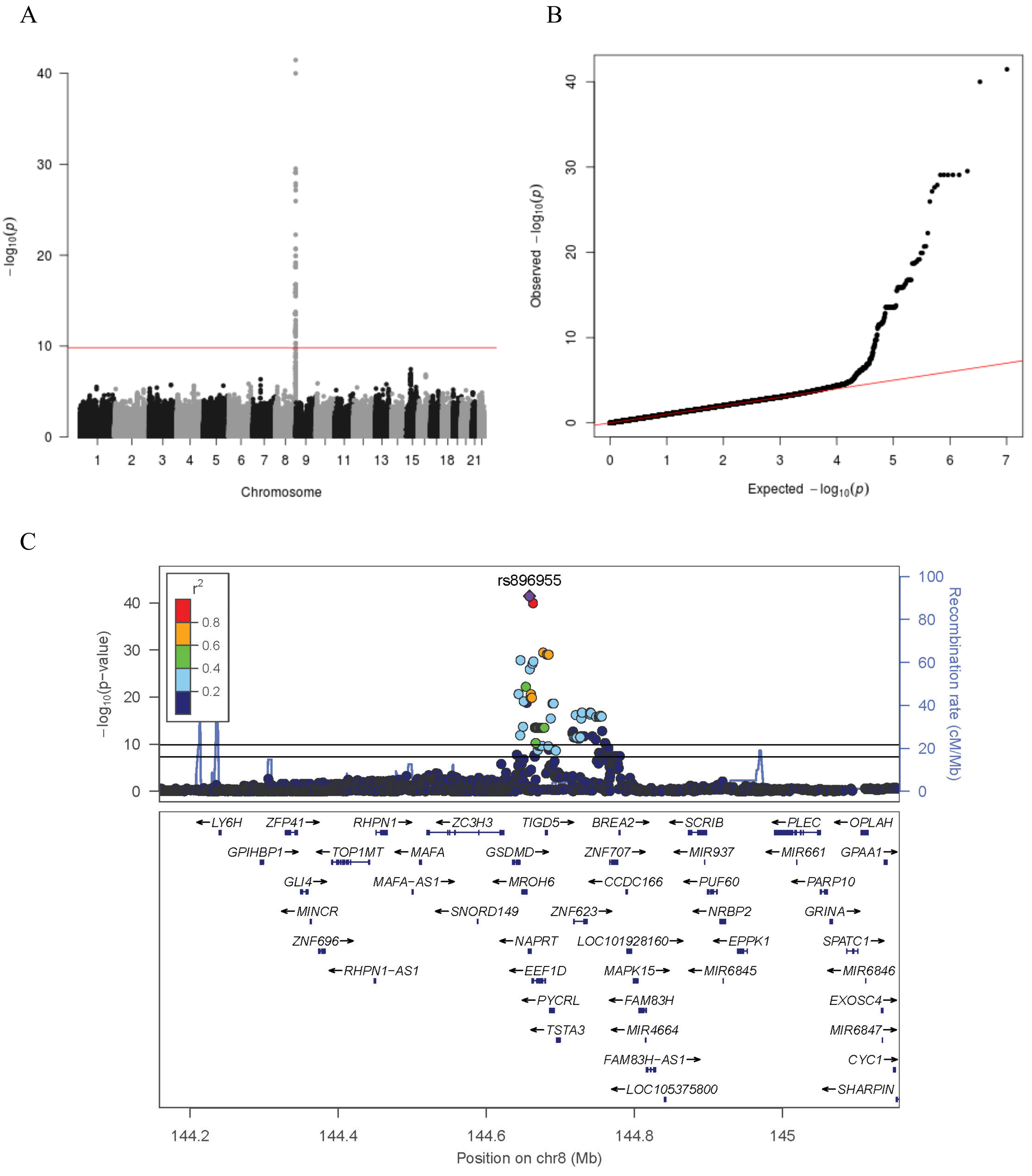

**Supplementary Figure 10. GWAS meta-analysis results for 1-ribosyl-imidazoleacetate (X61868)**

Panel A shows the Manhattan plot, with a red line indicating the Bonferroni-corrected significance threshold (P = $1.48\text{x}10^{-10}$). Panel B shows the Q-Q plot of the estimated effect sizes. Panel C shows the regional association plot for the associated locus. The top horizontal line represents the Bonferroni-corrected p-value threshold while the bottom line represents the standard genome-wide significance threshold.

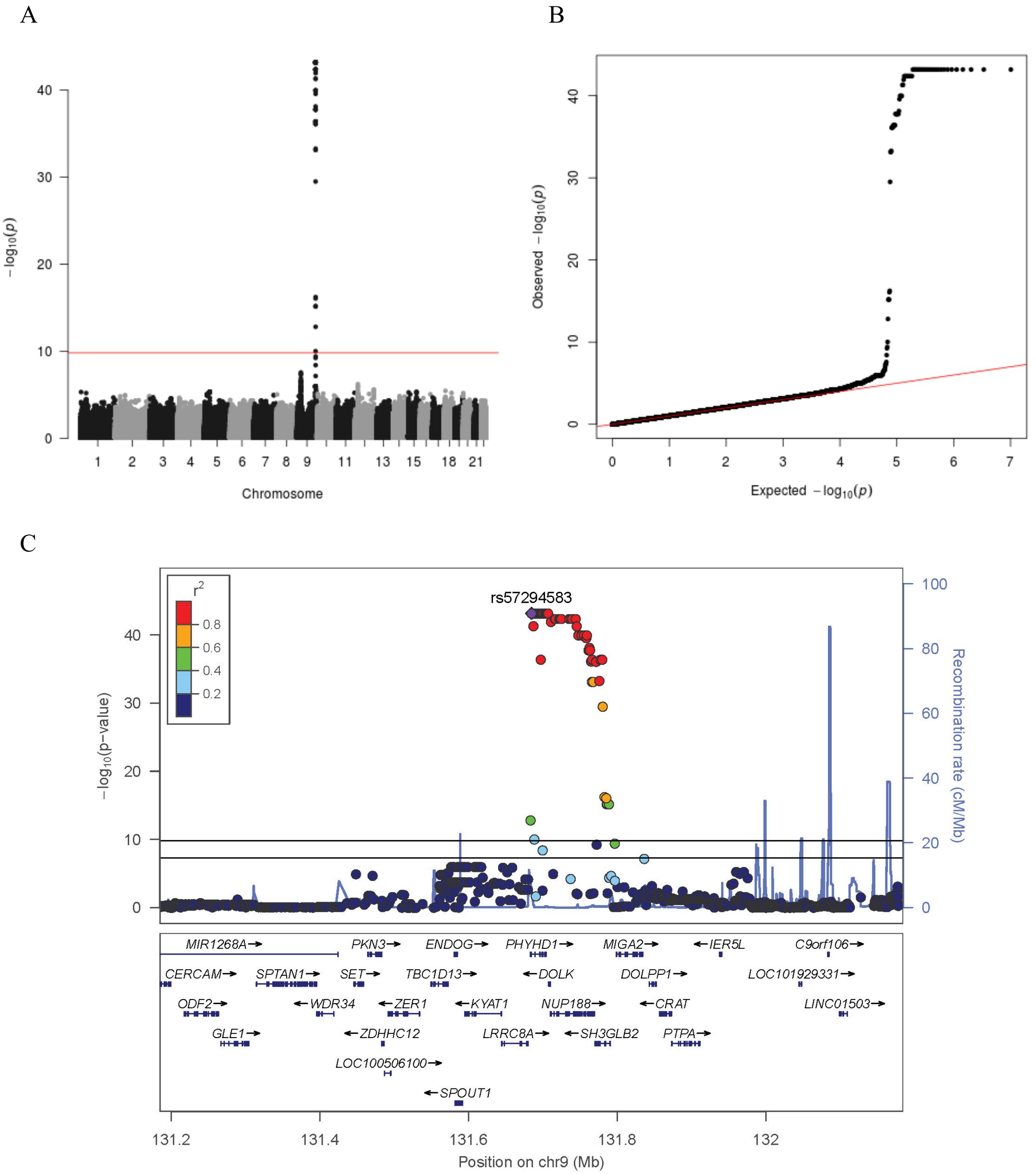

**Supplementary Figure 11. GWAS meta-analysis results for 2'-O-methylcytidine (X57554)**

Panel A shows the Manhattan plot, with a red line indicating the Bonferroni-corrected significance threshold ($P = 1.48 \times 10^{-10}$). Panel B shows the Q-Q plot of the estimated effect sizes. Panel C shows the regional association plot for the associated locus. The top horizontal line represents the Bonferroni-corrected p-value threshold while the bottom line represents the standard genome-wide significance threshold.

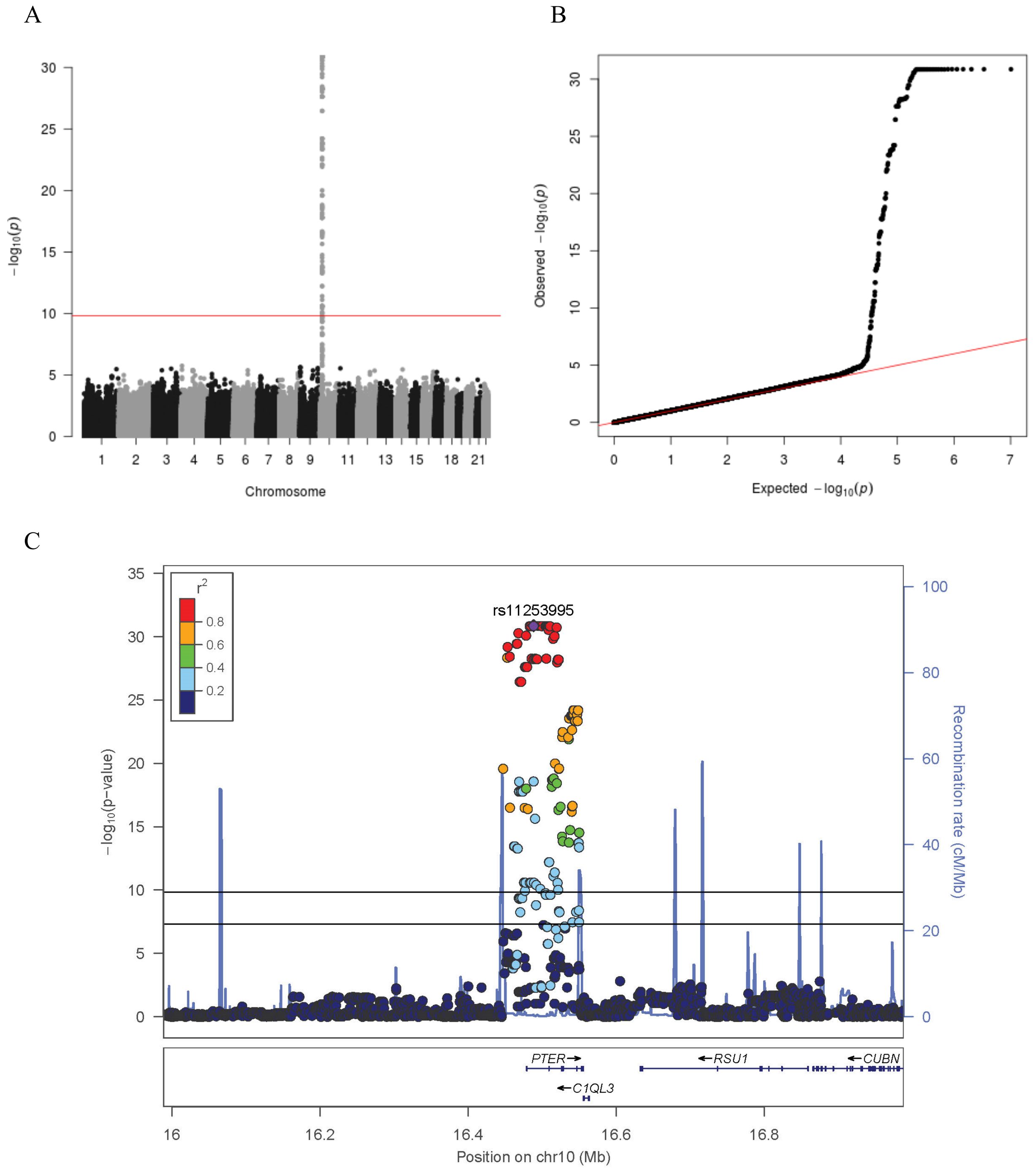

**Supplementary Figure 12. GWAS meta-analysis results for N-acetyl-beta-alanine (X37432)**

Panel A shows the Manhattan plot, with a red line indicating the Bonferroni-corrected significance threshold (P = $1.48x10^{-10}$). Panel B shows the Q-Q plot of the estimated effect sizes. Panel C shows the regional association plot for the associated locus. The top horizontal line represents the Bonferroni-corrected p-value threshold while the bottom line represents the standard genome-wide significance threshold.

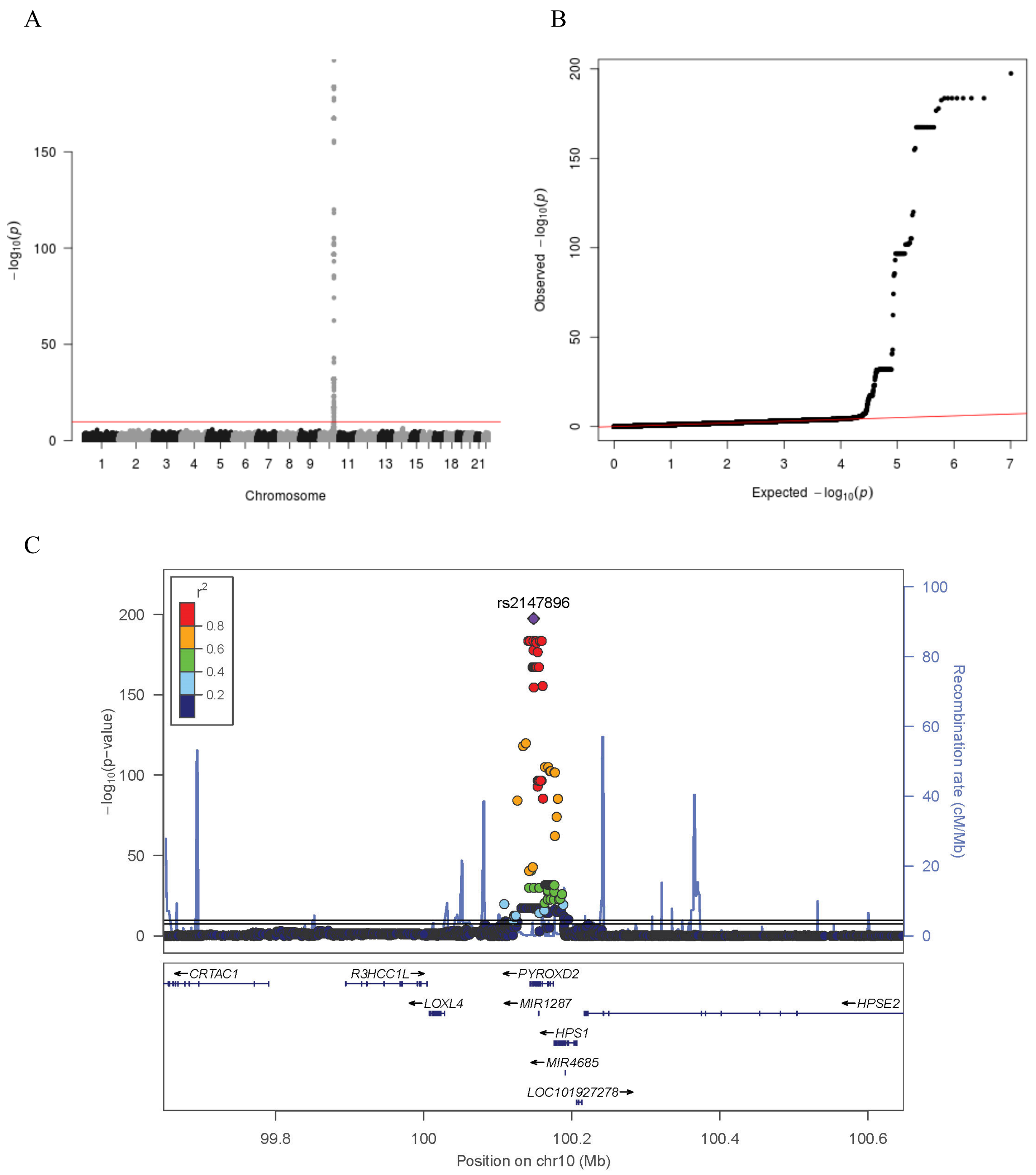

**Supplementary Figure 13. GWAS meta-analysis results for N6-methyllysine (X62249)**

Panel A shows the Manhattan plot, with a red line indicating the Bonferroni-corrected significance threshold ($P = 1.48x10^{-10}$). Panel B shows the Q-Q plot of the estimated effect sizes. Panel C shows the regional association plot for the associated locus. The top horizontal line represents the Bonferroni-corrected p-value threshold while the bottom line represents the standard genome-wide significance threshold.

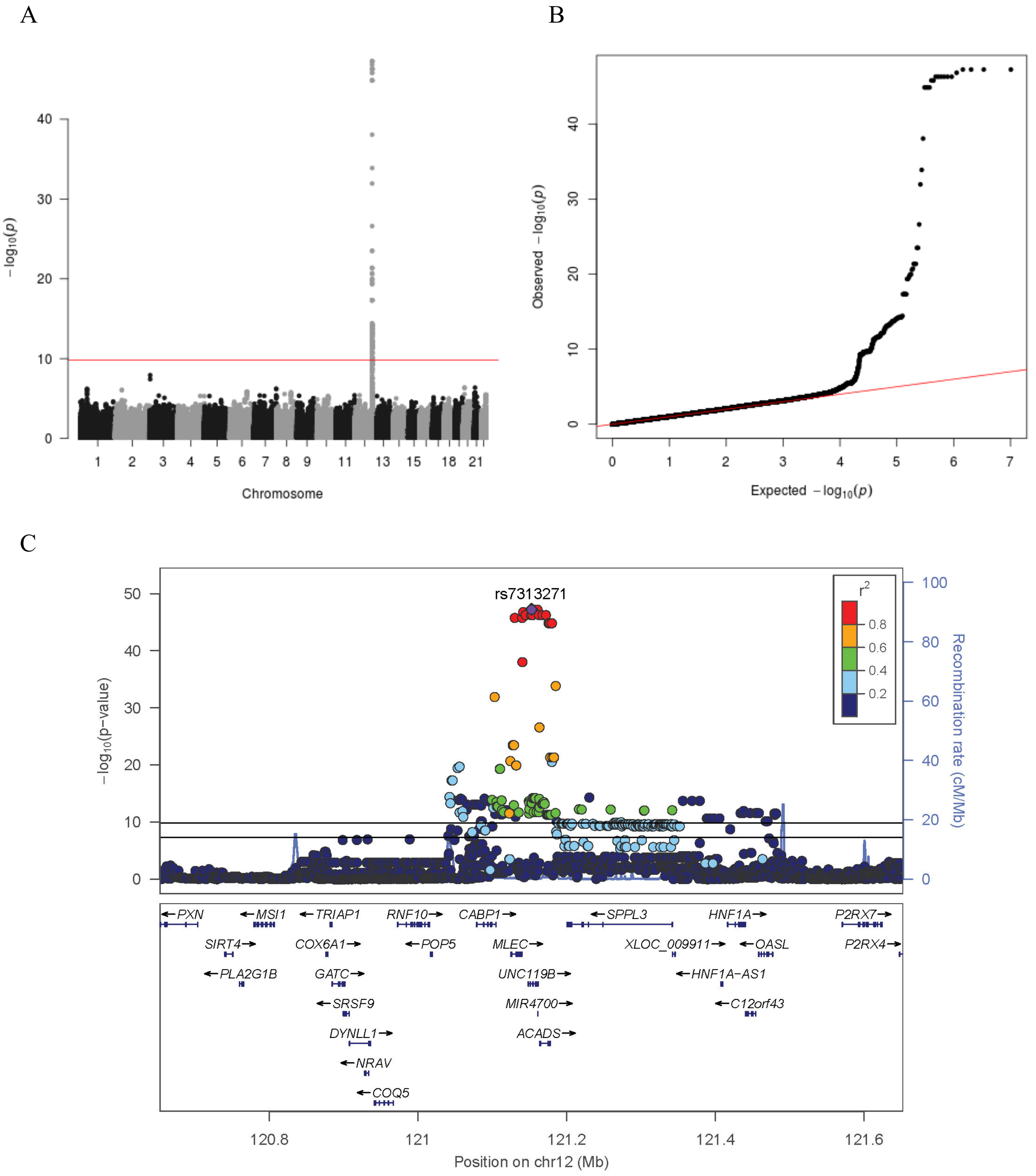

**Supplementary Figure 14. GWAS meta-analysis results for ethylmalonate (X15765)**

Panel A shows the Manhattan plot, with a red line indicating the Bonferroni-corrected significance threshold (P = $1.48 \times 10^{-10}$). Panel B shows the Q-Q plot of the estimated effect sizes. Panel C shows the regional association plot for the associated locus. The top horizontal line represents the Bonferroni-corrected p-value threshold while the bottom line represents the standard genome-wide significance threshold.

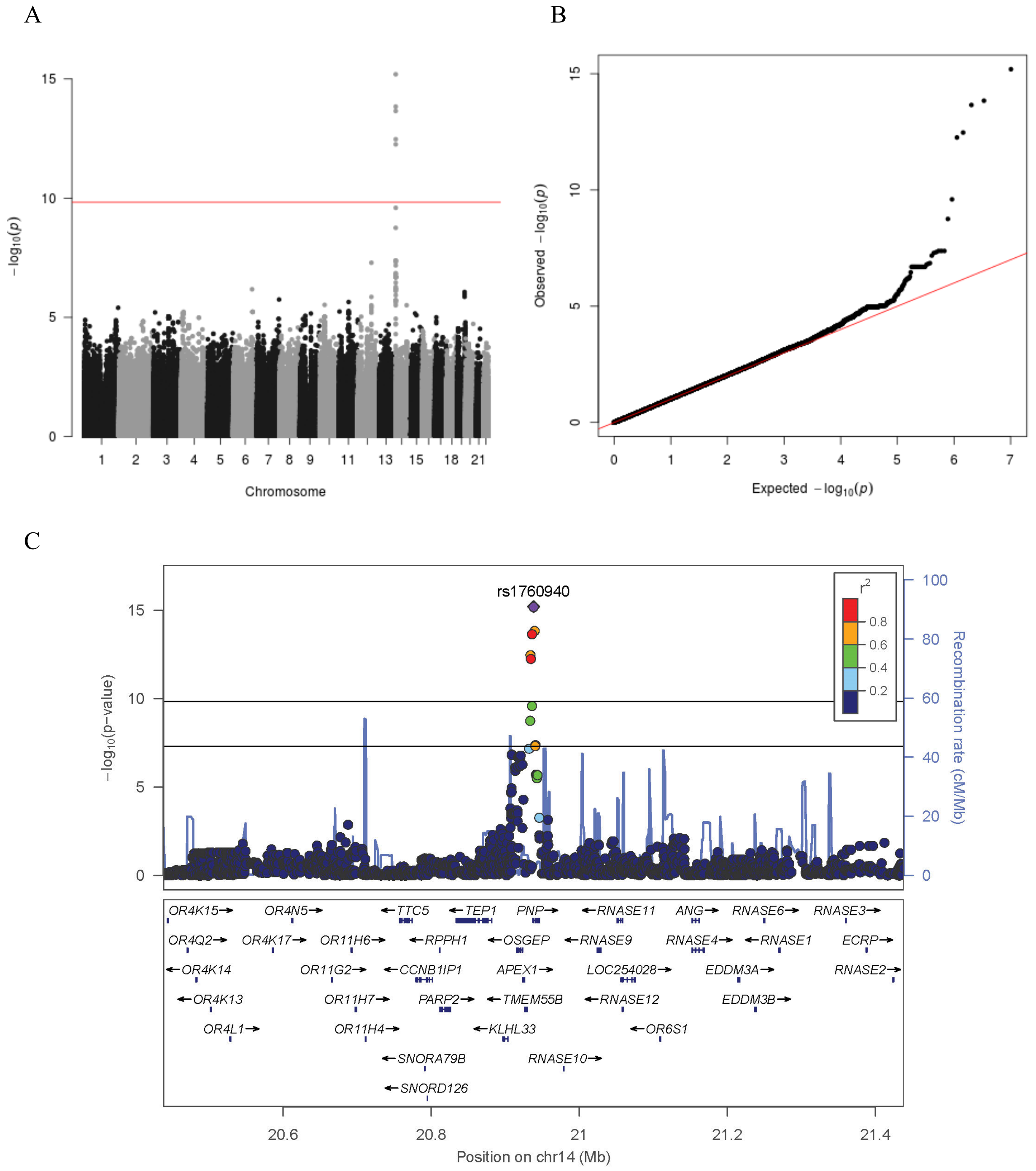

**Supplementary Figure 15. GWAS meta-analysis results for guanosine (X1573)**

Panel A shows the Manhattan plot, with a red line indicating the Bonferroni-corrected significance threshold (P = $1.48x10^{-10}$). Panel B shows the Q-Q plot of the estimated effect sizes. Panel C shows the regional association plot for the associated locus. The top horizontal line represents the Bonferroni-corrected p-value threshold while the bottom line represents the standard genome-wide significance threshold.

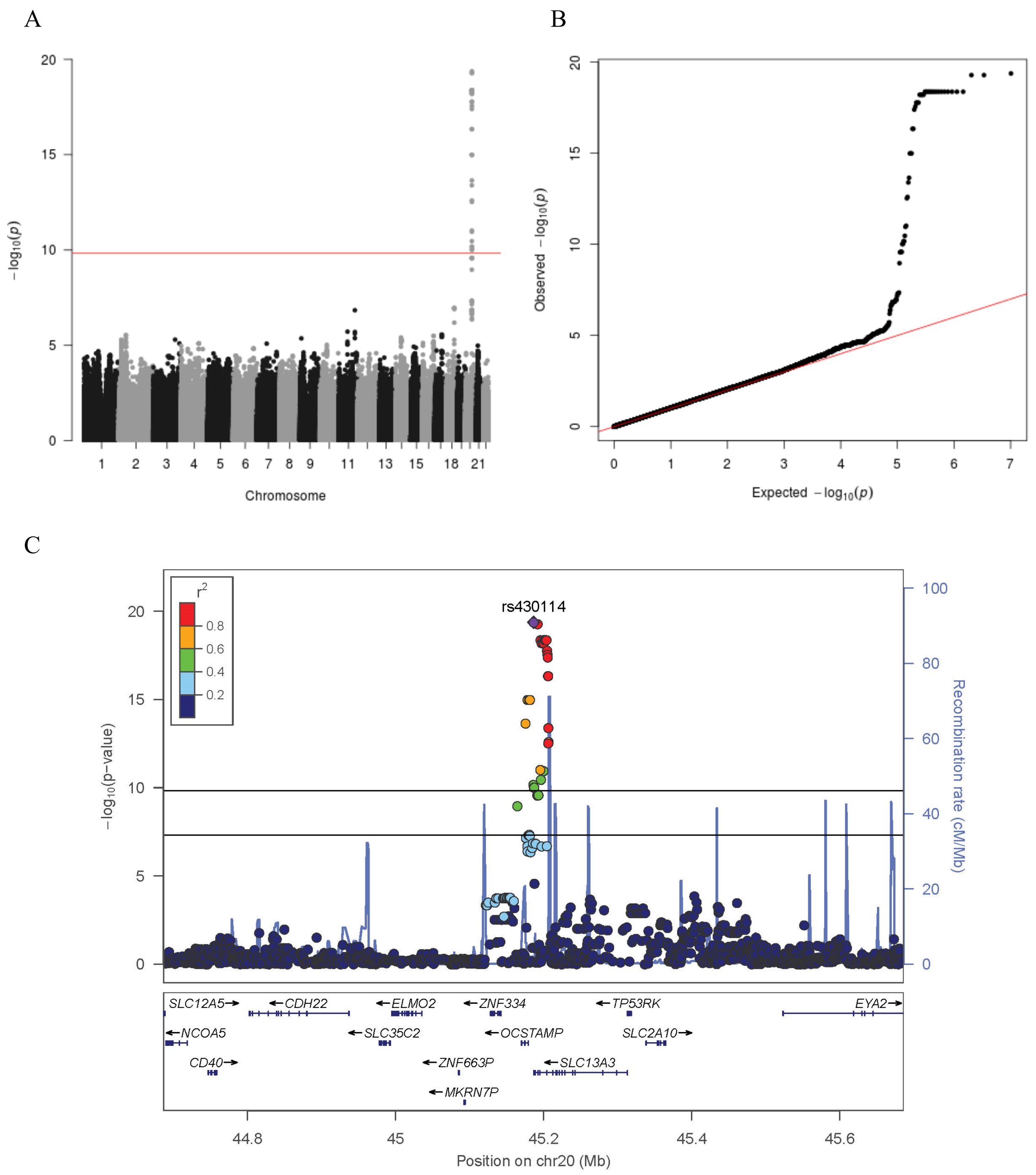

**Supplementary Figure 16. GWAS meta-analysis results for N-acetylglutamate (X15720)**

Panel A shows the Manhattan plot, with a red line indicating the Bonferroni-corrected significance threshold ($P = 1.48 \times 10^{-10}$). Panel B shows the Q-Q plot of the estimated effect sizes. Panel C shows the regional association plot for the associated locus. The top horizontal line represents the Bonferroni-corrected p-value threshold while the bottom line represents the standard genome-wide significance threshold.

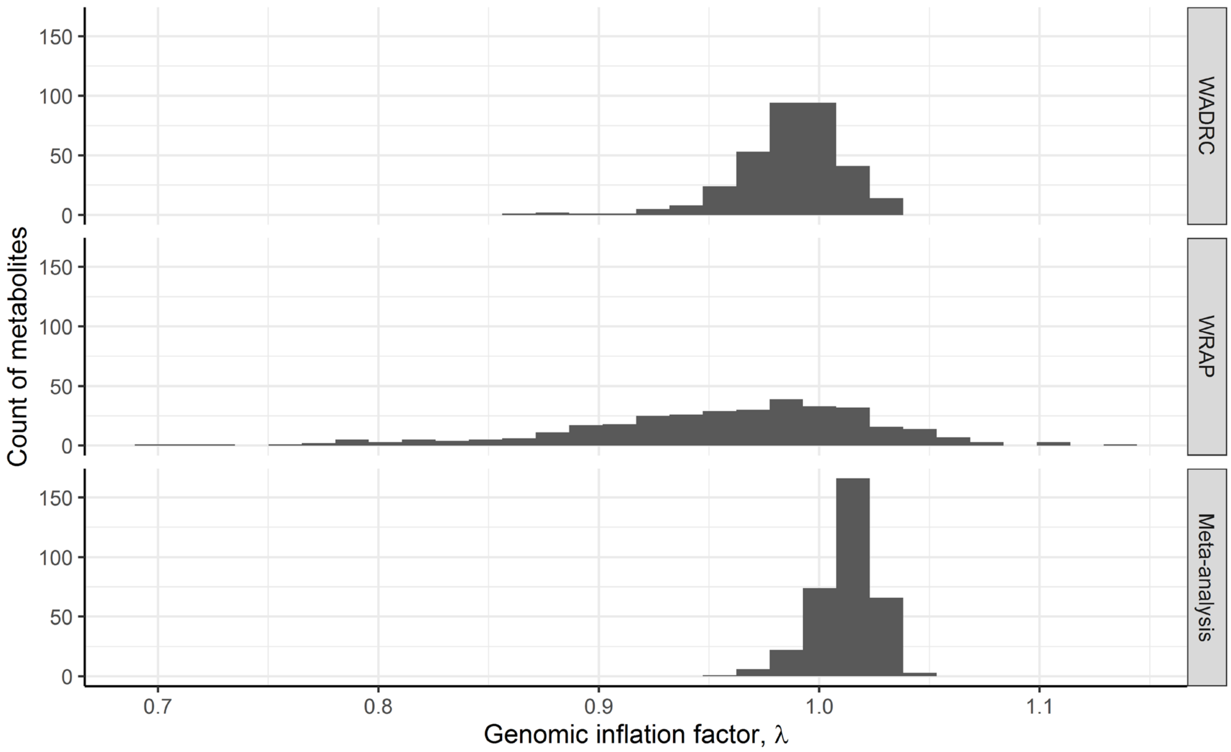

**Supplementary Figure 17. Genomic inflation factor distribution**

The distribution of the genomic inflation factors from the set of GWAS performed in the discovery (WADRC), replication (WRAP), and meta-analysis analyses is shown above.

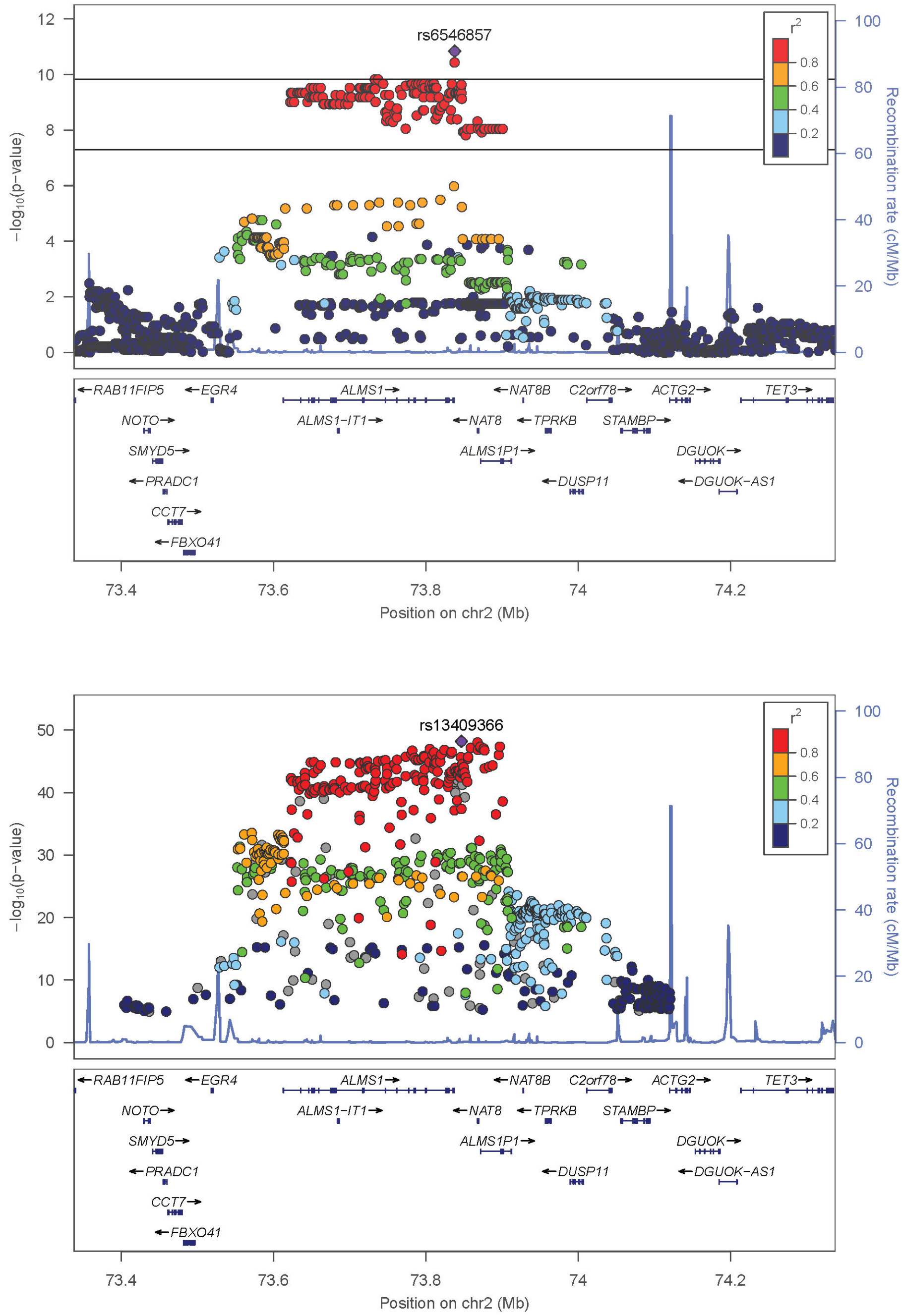

**Supplementary Figure 18. Side-by-side regional association plots for methionine sulfone (X44878)**

The regional association plots from CSF (top) and blood (bottom[1]). In the plot for CSF, the top horizontal line represents the Bonferroni-corrected p-value threshold, while the bottom line represents the standard genome-wide significance threshold. Non-CSF plots were generated manually using publicly available summary statistics.

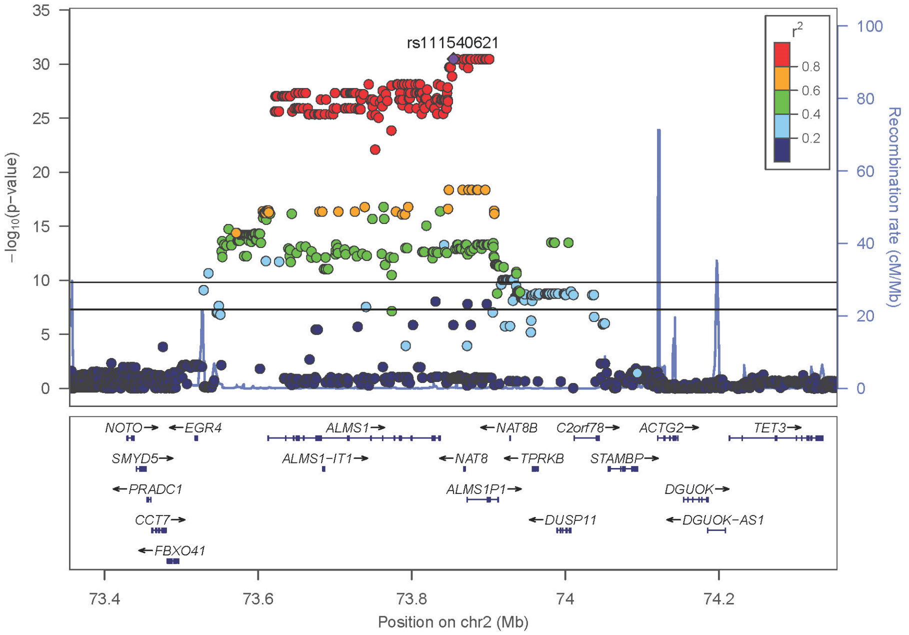

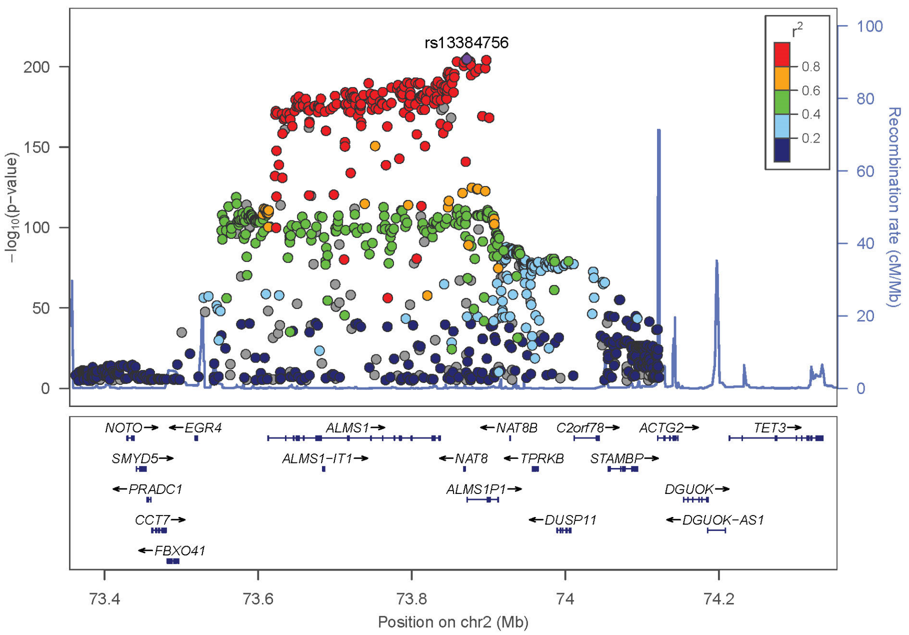

**Supplementary Figure 19. Side-by-side regional association plots for N-delta-acetylornithine (X43249)**

The regional association plots from CSF (top) and blood (bottom[1]). In the plot for CSF, the top horizontal line represents the Bonferroni-corrected p-value threshold, while the bottom line represents the standard genome-wide significance threshold. Non-CSF plots were generated manually using publicly available summary statistics.

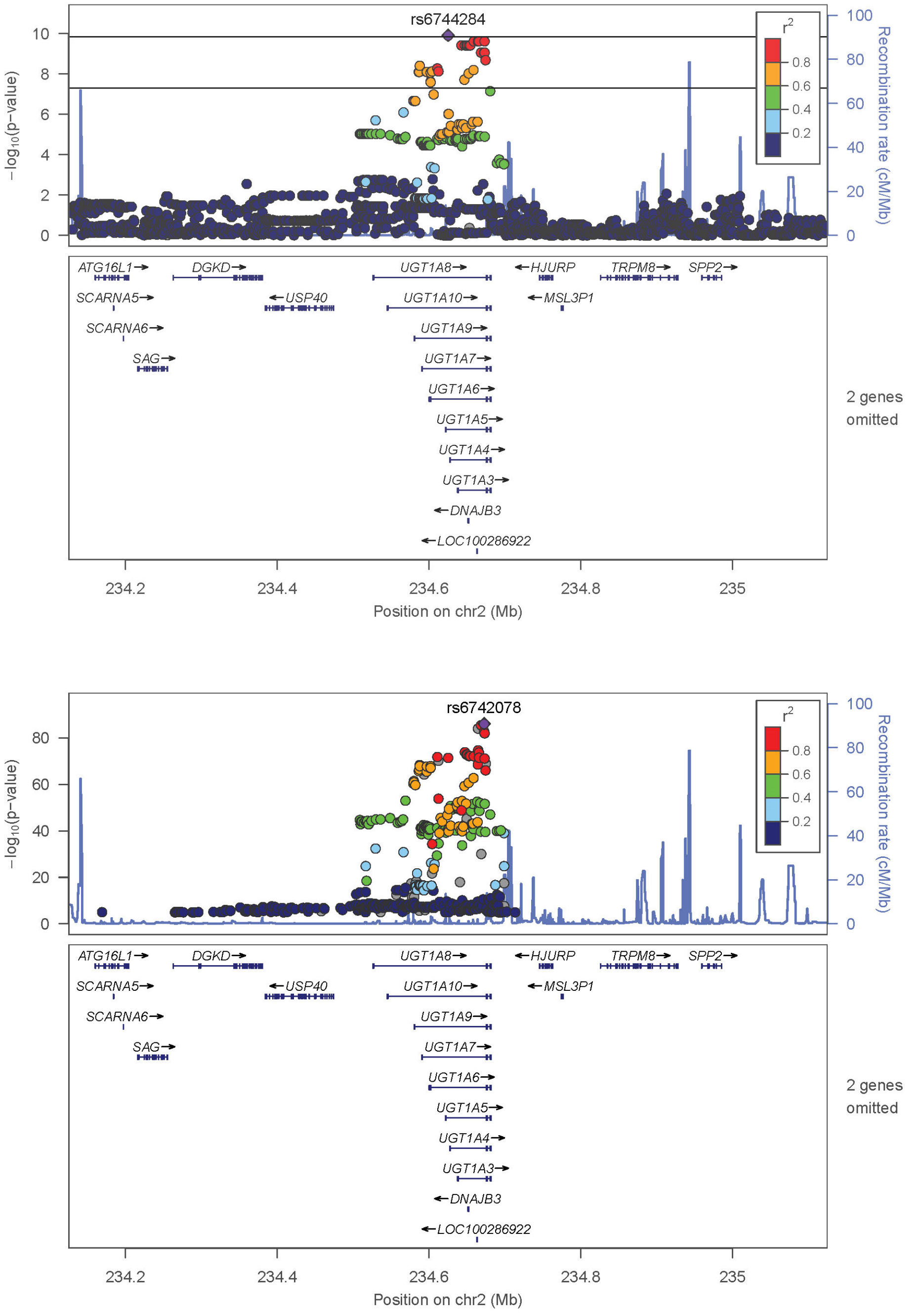

**Supplementary Figure 20. Side-by-side regional association plots for bilirubin (X43807)**

The regional association plots from CSF (top) and blood (bottom[1]). In the plot for CSF, the top horizontal line represents the Bonferroni-corrected p-value threshold, while the bottom line represents the standard genome-wide significance threshold. Non-CSF plots were generated manually using publicly available summary statistics.

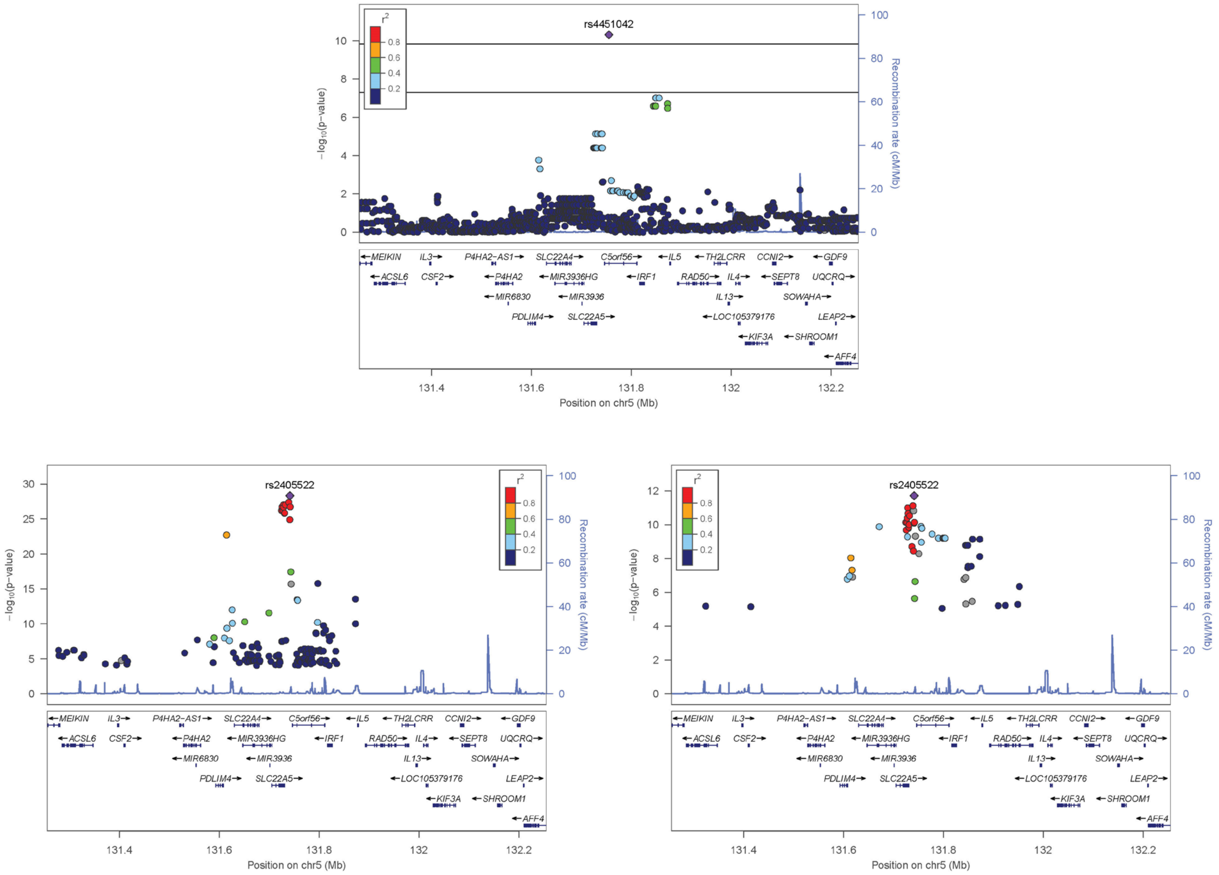

**Supplementary Figure 21. Side-by-side regional association plots for tryptophan betaine (X37097)**

The regional association plots from CSF (top) and blood (bottom-left[2] and bottom-right[1]). In the plot for CSF, the top horizontal line represents the Bonferroni-corrected p-value threshold, while the bottom line represents the standard genome-wide significance threshold. Non-CSF plots were generated manually using publicly available summary statistics.

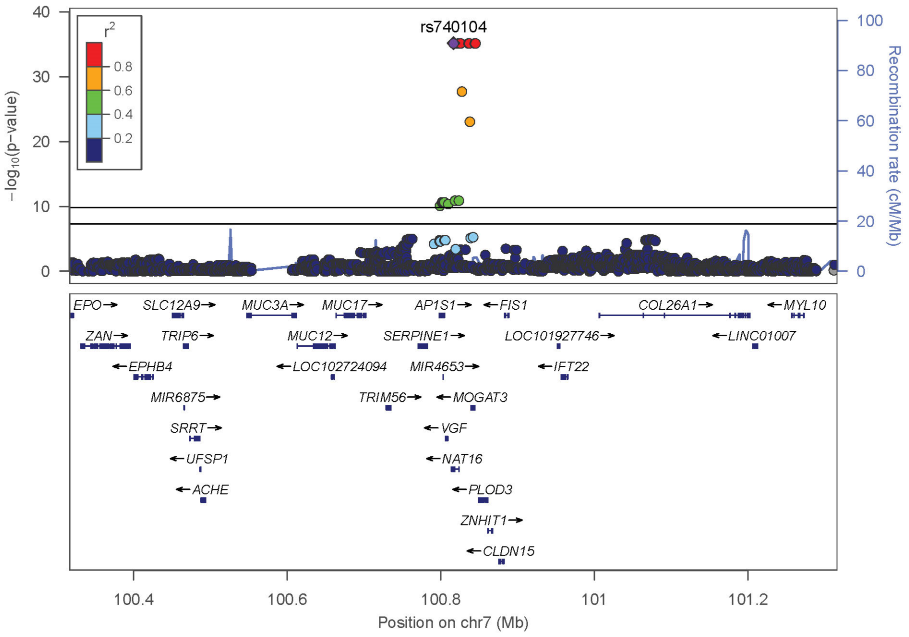

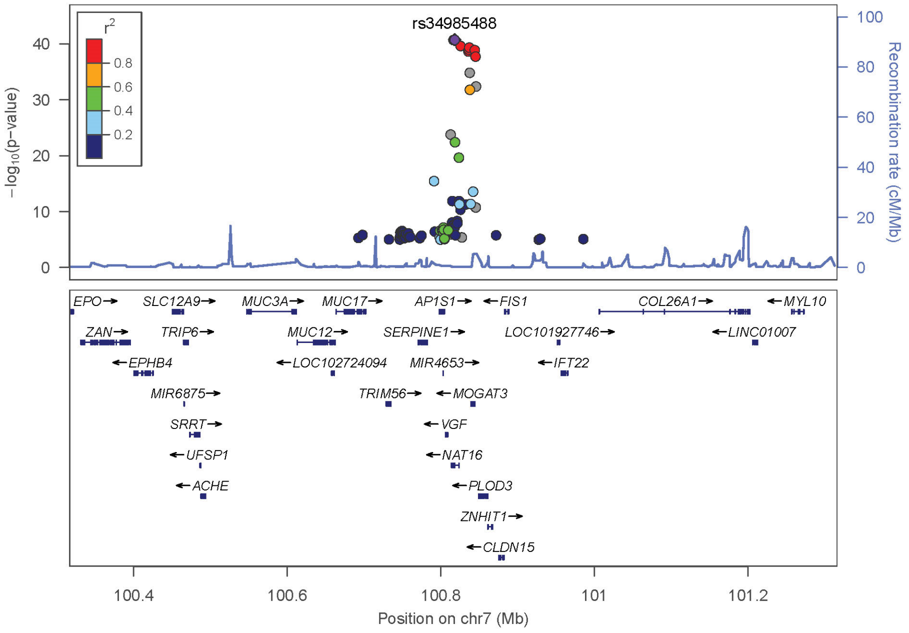

**Supplementary Figure 22. Side-by-side regional association plots for N-acetylhistidine (X33946)**

The regional association plots from CSF (top) and blood (bottom[1]). In the plot for CSF, the top horizontal line represents the Bonferroni-corrected p-value threshold, while the bottom line represents the standard genome-wide significance threshold. Non-CSF plots were generated manually using publicly available summary statistics.

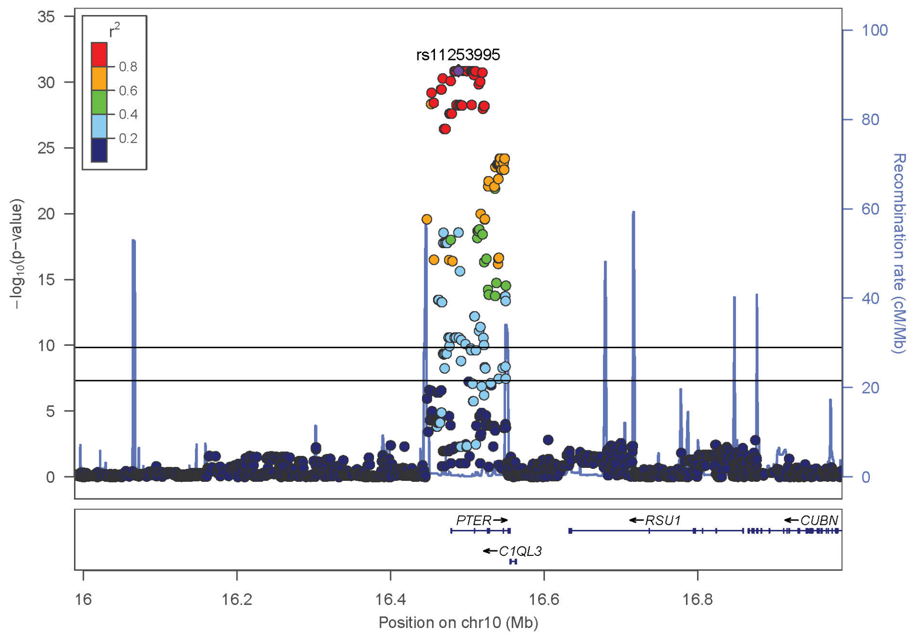

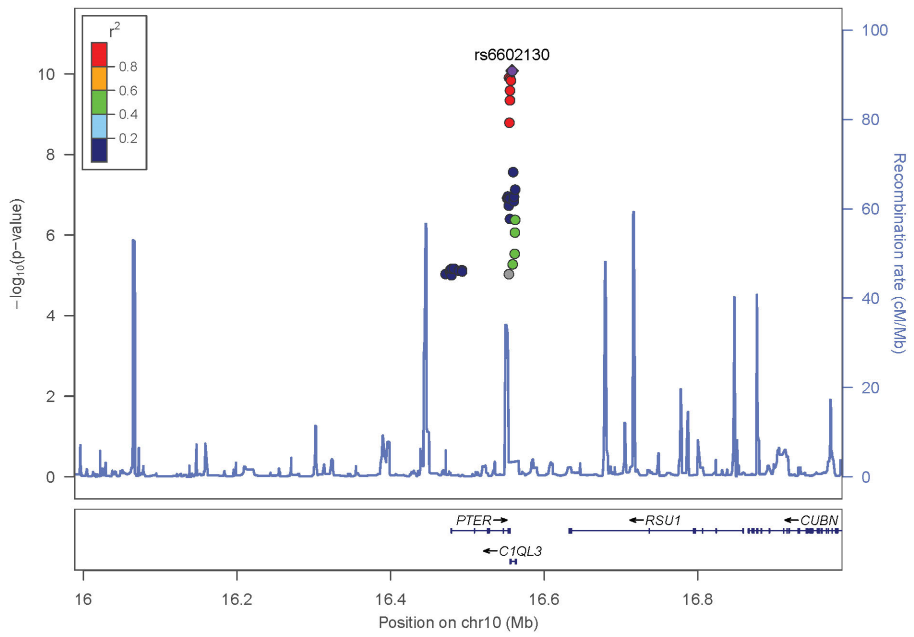

**Supplementary Figure 23. Side-by-side regional association plots for N-acetyl-beta-alanine (X37432)**

The regional association plots from CSF (top) and blood (bottom[1]). In the plot for CSF, the top horizontal line represents the Bonferroni-corrected p-value threshold, while the bottom line represents the standard genome-wide significance threshold. Non-CSF plots were generated manually using publicly available summary statistics.

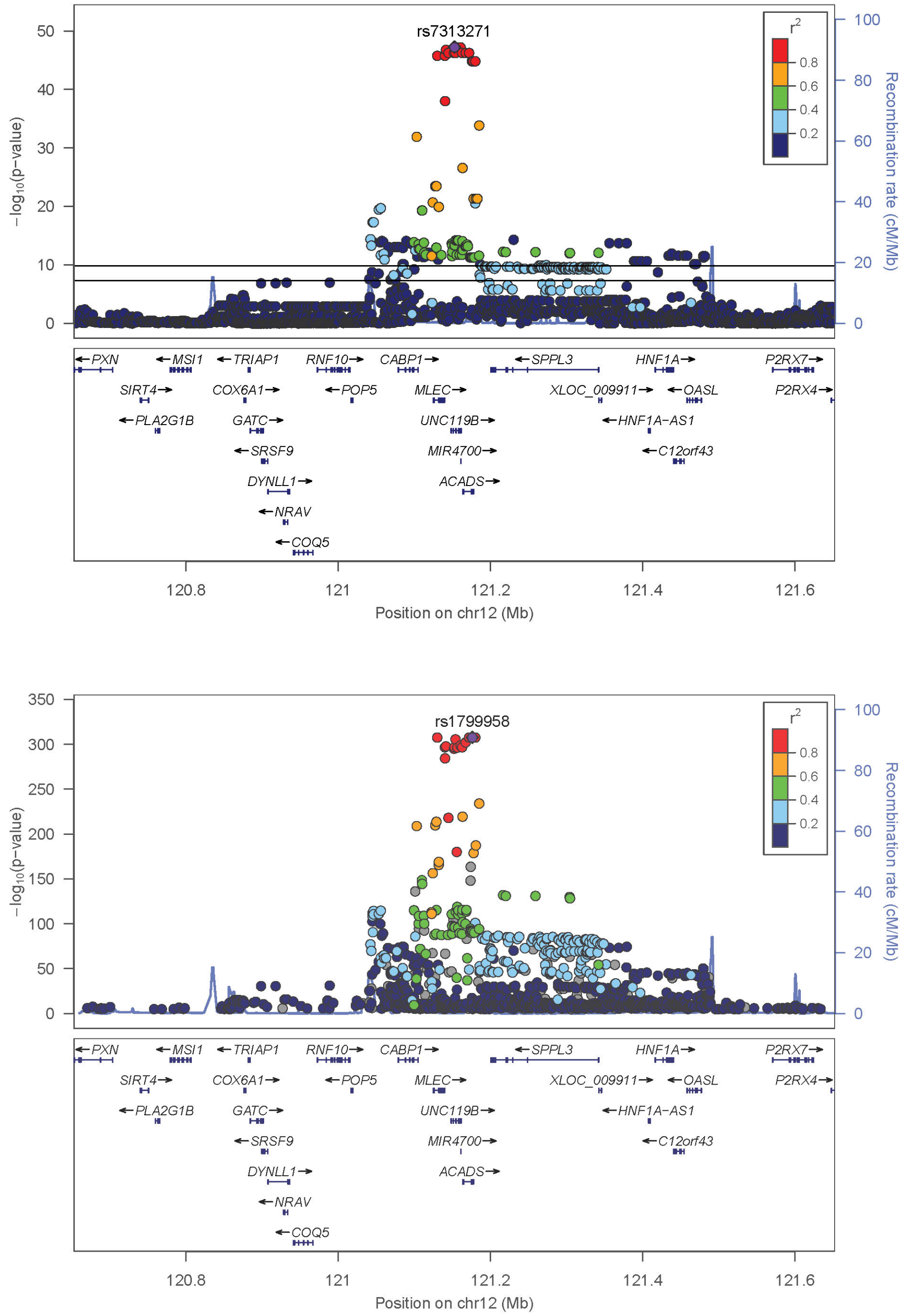

**Supplementary Figure 24. Side-by-side regional association plots for ethylmalonate (X15765)**

The regional association plots from CSF (top) and blood (bottom[1]). In the plot for CSF, the top horizontal line represents the Bonferroni-corrected p-value threshold, while the bottom line represents the standard genome-wide significance threshold. Non-CSF plots were generated manually using publicly available summary statistics.

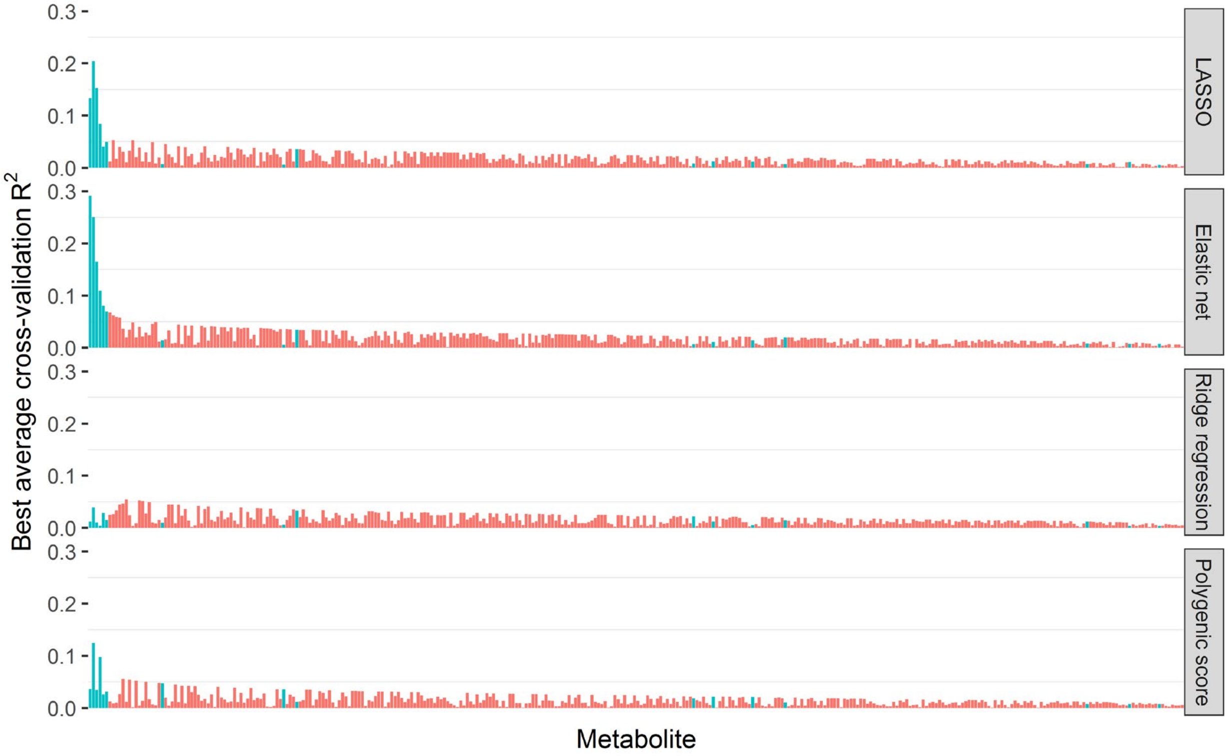

**Supplementary Figure 25. Metabolite prediction model performance by model type**

The prediction performance of the best model of each model type is shown, with metabolites arranged left-to-right according to the best possible model correlation across all models. Metabolites with a significant locus from the GWAS meta-analysis are highlighted in blue.

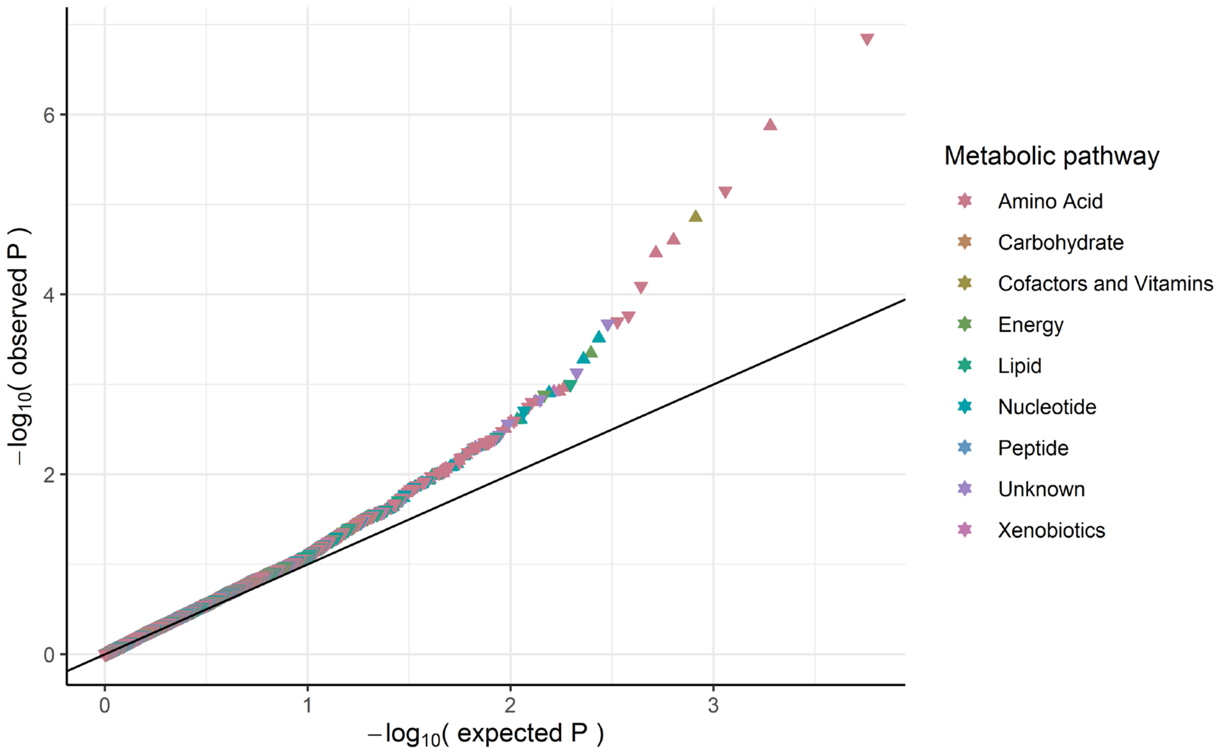

**Supplementary Figure 26. Metabolite-phenotype association analysis Q-Q plot**

Each point represents the association of one metabolite with a psychiatric or neurological phenotype, with the color representing the metabolite's metabolic pathway. Upward pointing triangles designate a positive association of the metabolite with the phenotype; downward pointing triangles designate a negative association.
